# Three Plasmid Strategies, One Intermediate Convergence State: Lineage-Specific Resistance–Virulence Architecture in Dominant Indian Carbapenem-Resistant *Klebsiella pneumoniae* Clones

**DOI:** 10.64898/2026.06.23.734134

**Authors:** Sanika Mahesh Kulkarni, Jobin John Jacob, Subbulaksmi Rajendra, Preethi S, Monisha Priya T, Aravind Velmurugan, Rosemol Nelson, Ayyanraj Neeravi, Lavanya Balaji, Karthik Gunasekaran, Abi Manesh, Rajni Ekadasi, Kamini Walia, Balaji Veeraraghavan

## Abstract

Carbapenem-resistant *Klebsiella pneumoniae* (CRKp) is a critical global healthcare threat driven by high-risk multidrug-resistant (MDR) clones that acquire hypervirulence genes. Although resistance-virulence co-occurrence is extensively documented, the plasmid-level mechanisms facilitating this convergence remain unclear. In this study, we utilized hybrid short- and long-read whole-genome sequencing of 376 clinical CRKp strains to define the evolutionary trajectories and structural plasmid dynamics of three predominant high-risk clones: ST147 (*n*=157), ST231 (*n*=108), and ST2096 (*n*=111). Carbapenemase genes were present in 90% of isolates, predominantly *bla*_OXA-48-like_ and *bla*_NDM-5_ co-harbored with *bla*_CTX-M-15_. Virulence profiling indicated high aerobactin (*iuc*) prevalence (62.7%), while salmochelin and colibactin were undetected. Hypermucoviscosity occurred infrequently (6.6%) and was independent of *rmpA*/*rmpA2*, confirming a clear genotype-phenotype discordance. Comparative plasmid mapping revealed three distinct, lineage-specific plasmid configurations underlying this intermediate convergent pathotype: ST147 exhibited dynamic, mosaic hybrid *IncFIB*-*IncHI1B* plasmids; ST2096 showed structurally stabilized hybrids; and ST231 retained virulence and resistance determinants on separate, segregated plasmids. These findings show that convergence is regulated by multiple, clone-specific evolutionary routes rather than a single path, highlighting the critical need for more in-depth genomic surveillance capable of identifying convergent plasmids along with high-risk lineages

## 1. Introduction

*Klebsiella pneumoniae* (Kp) is a major cause of hospital-acquired infections globally, with several clonal lineages demonstrating an exceptional capacity to acquire and disseminate antimicrobial resistance (AMR) and virulence determinants (1–3). The global epidemiology of epidemic *K. pneumoniae* is characterised by distinct geographic heterogeneity with ST258 predominates in North America and Europe, ST11 (clonal group 258) dominates in East Asia, while ST147 has emerged as a globally disseminated lineage associated with carbapenem resistance (7,8,9,12,13,22). Additional clones, including ST15 (13) and ST307 (12, 23), have achieved regional or international prominence, although none challenge the global dominance of ST258, ST11 (23), and ST147 (8,12,25). Consistent with this population structure, carbapenem resistance mechanisms also vary geographically: ST258 is predominantly associated with KPC carbapenemases in Europe and North America; ST11 in East Asia commonly carries KPC and/or NDM; while lineages circulating in South Asia and the Middle East are strongly associated with NDM and OXA-48–like carbapenemases (11,14,15,19,22).

Beyond the expansion of classical multidrug-resistant (MDR) lineages, increasing concern has focused on the emergence of *K. pneumoniae* strains in which AMR and hypervirulence traits converge within a single genetic background (16,20,24). Historically, MDR and hypervirulent *K. pneumoniae* (hvKp) populations were largely distinct; however, genomic investigations increasingly document convergence driven either by acquisition of virulence loci by high-risk resistant clones or by introduction of resistance determinants into traditionally hypervirulent lineages (24,26,28). Resistance acquisition by hvKp, particularly clonal group 23 (CG23; ST23) has been increasingly reported from Europe and North America. Conversely, virulence acquisition has been documented in several globally disseminated MDR lineages, including ST11 and ST147, most commonly mediated by horizontal transfer of loci encoding key determinants such as *rmpA*/*rmpA2,* aerobactin (*iuc*), and salmochelin (*iro*) (5,7,16,18,20). ST11 has emerged as a major driver of such convergence events in East Asia, where carbapenem-resistant hypervirulent *K. pneumoniae* (CR-hvKp) has caused large hospital outbreaks associated with high mortality. Similar convergence events are increasingly reported from South and Southeast Asia, highlighting the global expansion of this high-risk evolutionary strategy (15,27).

In South Asia, and particularly in India, genomic epidemiology studies reveal a distinct hospital-associated *K. pneumoniae* population structure dominated by high-risk clones including ST231, ST147, and the emerging lineage ST2096 (27,29). These lineages account for a substantial proportion of carbapenem-resistant infections in healthcare settings and represent the principal genetic backgrounds in which resistance–virulence convergence has been observed. Notably, convergence among Indian isolates differs from classical descriptions involving acquisition of complete pLVPK-like virulence plasmids. Instead, it is predominantly characterised by acquisition of discrete virulence loci most frequently *iuc*, alone or in combination with *rmpA, rmpA2*, or *iro* often embedded within hybrid plasmids that also carry key AMR genes, including carbapenemases (27,29,30). Such hybrid resistance–virulence plasmids may provide a stable mechanism for maintaining convergent phenotypes in hospital settings.

Recent refinements to the definition of hypervirulent *K. pneumoniae* emphasise the presence of at least four of five key virulence markers (*rmpA, rmpA2, iucA, iroB*, and peg-344). Under this framework, convergent strains circulating among dominant Indian high-risk clones (ST231, ST147, and ST2096) typically do not meet criteria for multidrug-resistant hypervirulent *K. pneumoniae* (MDR-hvKp), as they harbour only a subset of these loci. Nevertheless, accumulating evidence suggests that such partially virulent strains may represent an intermediate pathogenic state along a virulence continuum between classical *K. pneumoniae* (cKp) and hvKp (1,24,29,30). In particular, iuc-positive strains lacking the full complement of hypervirulence markers have been proposed to occupy a transitional state with important clinical and epidemiological implications (24).

In this study, we present the most comprehensive plasmid-resolved genomic characterisation to date of the three dominant carbapenem-resistant *K. pneumoniae* clones circulating in Indian hospitals ST147, ST231, and ST2096. By integrating long-read hybrid genome assembly with core-genome phylogenetics and comprehensive plasmid structural analysis across 376 genomes spanning three years of prospective surveillance and archived isolates, we aimed to elucidate the distinct lineage-specific plasmid strategies underlying resistance–virulence convergence, to characterise the structural diversity of hybrid resistance–virulence plasmids, and to define the spectrum of intermediate resistance–virulence convergence states across major hospital-associated *K. pneumoniae* lineages in India.

## 2. Methodology

### 2.1 Study design and clinical isolates

This retrospective study was conducted between January 2023 and December 2025 at Christian Medical College (CMC), Vellore, India. All clinical *K. pneumoniae* isolates recovered from cases of bloodstream infection (BSI) and pneumonia during the study period were included in the primary surveillance dataset. For the present genomic analysis, a subset of isolates (n=271) was selected based on multilocus sequence typing (MLST), focusing on the dominant high-risk clones ST147, ST231, and ST2096, for which whole-genome sequencing data were available as part of the primary study. To provide historical context and assess temporal changes in antimicrobial resistance and virulence, additional genomes belonging to these high-risk clones and collected prior to the study period (n=105) were also included for comparative analysis. All patient-identifiable information was removed prior to analysis to ensure confidentiality. The study was approved by the Institutional Review Board and Ethics Committee of Christian Medical College, Vellore (IRB Min. No. 15248 dated 22.03.2023). A complete list of the isolates and associated metadata is provided in the Supplementary Table 1. All archived isolates were revived on MacConkey agar for further phenotypic and molecular testing.

### 2.2 Phenotypic characterization

The hypermucoviscosity (hmv) phenotype was evaluated using the string test (43). Briefly, isolates were cultured overnight on MacConkey agar, and a sterile inoculating loop was used to check the string; the formation of a viscous string measuring >5 mm was interpreted as a positive result.

Antimicrobial susceptibility testing (AST) was performed on 271 isolates using the Kirby–Bauer disk diffusion method, and results were interpreted according to CLSI guidelines (M100-S35). Inhibition zone diameters were measured and interpreted using CLSI-recommended breakpoints (CLSI, 2023–2025) for ertapenem, meropenem, piperacillin-tazobactam, cefoperazone-sulbactam, amoxicillin-clavulanate, cefepime, ceftazidime, amikacin, gentamicin, levofloxacin, and trimethoprim-sulfamethoxazole. *Escherichia coli* ATCC 25922 served as the quality control strain.

### 2.3 DNA Extraction and Detection of Virulence Genes

Genomic DNA was extracted directly from well-isolated colonies grown on MacConkey agar plates using the QIAamp blood mini kit (Qiagen,Hilden, Germany) following the instructions. The purity and concentration of extracted DNA were assessed using Nanodrop One (Thermo Scientific) and Qubit dsDNA HS Assay Kit (Life Technologies) respectively. Multiplex polymerase chain reaction (PCR) assays were performed to screen for key hypervirulence-associated genes, including *iucA*, *rmpA*, *rmpA2*, *iroB*, and *peg-344*, using previously described primers and protocols. In parallel, multiplex PCR assays were used to detect major extended-spectrum β-lactamase (ESBL) and carbapenemase genes, including *bla*_KPC_, *bla*_NDM_, *bla*_OXA-48–like_, and *bla*_CTX-M_ (4).

### 2.4 Whole Genome Sequencing and Assembly

Whole-genome sequencing (WGS) was performed for study isolates using short-read Illumina sequencing. Paired-end libraries were prepared with the Illumina kit and sequenced as 2 × 150 bp reads on an Illumina NovaSeq Xplus platform at Unipath Speciality Laboratory, Ahmedabad, India. For a subset of isolates, long-read sequencing was performed to enable complete genome and plasmid reconstruction. High-molecular-weight DNA libraries were prepared using the Oxford Nanopore Technologies (ONT) Rapid Sequencing DNA V14-barcoding Kit (SQK-RBK114.96) and sequenced on a PromethION P2 Solo device using (R10.4.1-FLO-PRO114M) flow cells with Q30+ chemistry, following the manufacturer’s protocol.

Quality assessment of Illumina short reads and Nanopore long reads was carried out using FastQC v0.11.9 (https://github.com/s-andrews/fastqc) and LongQC v1.2.0c, respectively. Illumina reads were trimmed and quality-filtered using Trimmomatic v0.39 (https://github.com/usadellab/trimmomatic) with a Phred quality score threshold of Q30. De novo assemblies from Illumina reads were generated using SKESA v2.4.0 (https://github.com/ncbi/SKESA). Hybrid genome assemblies combining Illumina and Nanopore reads were constructed using the Hybracter pipeline (https://github.com/gbouras13/hybracter).

### 2.5 Comparative Genome Analysis

Sequence types (STs) of all *K. pneumoniae* isolates were confirmed using the MLST tool (https://github.com/tseemann/mlst). Virulence-associated genes, including *rmpA*, *rmpA2*, *iucABCD*, *iroBCDN*, *peg-344*, *ybt*, and *clb*, were identified through Kleborate(v3.1.3) (https://github.com/katholt/Kleborate). Resistance genes and plasmid replicons were detected using ABRicate (https://github.com/tseemann/abricate) with the AMRFinderPlus (v4.2.5) and PlasmidFinder (v2.1.6) databases respectively. The capsular (K) and O antigen loci were typed using Kaptive (https://github.com/katholt/Kaptive). These analyses allowed for comparison of virulence gene profiles, capsular diversity, and antimicrobial resistance profiles across time and between lineages.

### 2.6 Phylogenetic Analysis

To provide a global perspective and ensure comprehensive lineage coverage, isolate IDs for *K. pneumoniae* ST147, ST231, and ST2096 were first identified using Pathogenwatch https://pathogen.watch/) and BIGSdb (https://bigsdb.pasteur.fr/klebsiella/) after which corresponding genome assemblies were retrieved from NCBI Pathogen detection (https://www.ncbi.nlm.nih.gov/pathogens/) or, where unavailable, raw sequencing reads were obtained from ENA and assembled in-house. An initial dataset was curated by retaining genomes with complete metadata (year, country, and isolation source) and acceptable assembly quality (N50 and contig number). A representative subset of genomes for each lineage was then selected using Treemmer (v0.3; https://github.com/niemasd/Treemmer) with the pruning option, ensuring proportional global sampling such that study isolates constituted at least 10% of each lineage-specific collection. Genomes showing evidence of contamination (e.g., mixed sequence types) or poor assembly quality were excluded following manual curation and quality control.

All genomes, including the global collection and study isolates, were annotated using Prokka (v1.14.6; https://github.com/tseemann/prokka). For the genetically diverse dataset, Prokka-annotated genome assemblies were used as input for Panaroo (v1.5.2) to perform pangenome reconstruction and define the collection core genome (https://github.com/gtonkinhill/panaroo). The resulting core genome alignment was subsequently used for phylogenetic inference. A maximum-likelihood core genome phylogeny was constructed using IQ-TREE v2.2.0 (https://github.com/iqtree/iqtree2), with the TN+F (Tamura–Nei with empirical base frequencies) substitution model selected as the best-fit model by ModelFinder, and nodal support evaluated using 10,000 bootstrap replicates. Based on the overall core genome phylogeny, isolates belonging to the ST147, ST231, and ST2096 lineages were identified and separately selected for further SNP-based phylogenetic analysis using Snippy (https://github.com/tseemann/snippy) to investigate lineage-specific genomic relatedness and population structure. Phylogenetic trees were based on custom reference constructed from the 629 conserved core genes defined in the Pasteur MLST scheme, retrieved from the Institut Pasteur BIGSdb database (https://bigsdb.pasteur.fr/) Core genome SNP alignments were used to reconstruct maximum-likelihood phylogenies using IQ-TREE v2.2.0, with the GTR+F+I+R10 (a General Time-Reversible model with empirical base frequencies, invariable sites, and 10 discrete gamma rate categories) identified as the best-fit substitution model using ModelFinder, and branch support assessed with 10,000 bootstrap replicates for all the lineage specific phylogenies. All phylogenetic trees were visualized and midpoint-rooted using iTOL (https://itol.embl.de/).

Clonal structure and population clustering were analysed using hierarchical Bayesian Analysis of Population Structure (hierBAPS) implemented via fastbaps (https://github.com/gtonkinhill/fastbaps), allowing hierarchical resolution at multiple levels of divergence (up to fifteen levels). Metadata such as capsular (K) loci, resistance and virulence gene profiles, and country of origin were mapped onto the trees to assess phylogeographic patterns and convergence events. Phylogeny was integrated with metadata, including K-locus type, resistance profile, and virulence profiles, to check for clonal expansion patterns. Comparative genomic analysis was performed to explore lineage-specific traits and plasmid backbones across ST231, ST2096, and ST147.

### 2.7 Plasmid Reconstruction and Structural Analysis

Hybrid-assembled genomes were used to reconstruct complete plasmid sequences. Plasmid replicon types were identified using MOB-suite (https://github.com/phac-nml/mob-suite) to infer plasmid mobility and to detect potential co-integrate or hybrid plasmid structures. Comparative structural analyses were performed to assess diversity, conservation, and rearrangements within plasmid backbones, with particular emphasis on regions carrying virulence- and antimicrobial resistance–associated loci. Special attention was given to the organization and integration of virulence determinants within plasmids that also harboured resistance genes, enabling characterization of resistance–virulence hybrid plasmid architectures. Hybrid-assembled plasmid sequences were visualized and manually inspected using Proksee (https://proksee.ca/) and EasyFig (https://mjsull.github.io/Easyfig/). Proksee was used for circular plasmid maps, while EasyFig was used for linear comparison of plasmid regions to identify conserved regions and structural rearrangements across representative plasmids.

## 3. Results

### 3.1 Clinical isolates, Lineage Distribution and Antimicrobial Susceptibility

A total of 700 *K. pneumoniae* isolates were recovered from blood and respiratory samples and sequenced at CMC Vellore between January 2023 and December 2025 as part of primary surveillance. For this study, we selected a subset of 271 isolates belonging to the high-risk clones based on their MLST profiles ST147, ST231, and ST2096, supplemented by 105 archived genomes of the same clones (total n=376), enabling assessment of temporal changes in antimicrobial resistance and virulence. The isolates within the three lineages were distributed as ST147 (n = 157), ST231 (n = 108), and ST2096 (n = 111). For phenotypic analysis, antimicrobial susceptibility testing was performed on 271 isolates, while archived isolates were included to provide historical context and support comparative genomic and temporal analyses.

Antimicrobial susceptibility testing confirmed uniformly low susceptibility across most antibiotic classes, reflecting the extensively drug-resistant phenotype characteristic of these lineages. Among carbapenems, susceptibility was 10.0% (27/271) to ertapenem and 10.7% (29/271) to meropenem, 4.8% (13/271) susceptible to amoxicillin/clavulanate, 5.5% (15/271) to piperacillin/tazobactam, and 8.1% (22/271) to cefoperazone/sulbactam. Third- and fourth-generation cephalosporins also showed low activity, with 8.1% (22/271) susceptibility to ceftazidime and 8.9% (24/271) to cefepime. Susceptibility to amikacin and gentamicin was 7.4% (20/271) and 8.9% (24/271), respectively, whereas comparatively higher susceptibility was observed for netilmicin (17.8%, 48/271) and tobramycin (18.1%, 49/271). Fluoroquinolone susceptibility was 19.3% (52/271) for ciprofloxacin and 8.9% (24/271) for levofloxacin. Cotrimoxazole susceptibility was detected in 7.4% (20/271) of isolates. Collectively, susceptibility to most first- and second-line agents was below 10%, underscoring the severely limited therapeutic options available for infections caused by these dominant lineages and highlighting the critical urgency of enhanced genomic surveillance.

### 3.2 Hypermucoviscosity phenotype

The hypermucoviscosity phenotype was infrequently observed among the study isolates. Only 18 of 271 isolates (6.6%) were positive by the string test, revealing a striking genotype–phenotype discordance: despite a high prevalence of virulence-associated loci particularly the aerobactin operon (62.7%) and *rmpA2* (41.8%) classical hypermucoviscosity remained uncommon across all three dominant lineages. This finding has important clinical implications: phenotypic screening by string test alone will substantially underestimate the true burden of virulence-competent isolates in these MDR lineages. It further confirms that in the context of Indian hospital-adapted clones, genomic detection of virulence determinants rather than phenotypic testing is the only reliable strategy for identifying convergent strains.

### 3.3 Genomic Characterization of Resistance Genes

Carbapenemase-encoding genes were detected in 90% (339/376) of genomes and frequently co-occurred with ESBL-associated genes (70%, 262/376), with 64.9% (244/376) harbouring both, indicating extensive convergence of β-lactam resistance mechanisms. The carbapenemase gene landscape was dominated by OXA-48–like variants, particularly *bla*_OXA-232_, detected either alone or in frequent combination with *bla*_NDM-5_. Dual-carbapenemase genotypes were common and predominantly involved *bla*_NDM-5_ and *bla*_OXA-48–like_ combinations, whereas *bla*_NDM-1_ was infrequent and *bla*_KPC_ was absent, defining a distinct carbapenemase profile.

ESBL-mediated resistance was largely driven by *bla*_CTX-M-15_, which was widespread across all lineages, consistent with stable maintenance across diverse genetic backgrounds. In addition to enzyme-mediated mechanisms, outer membrane porin alterations were nearly universal (99%), including truncations of *ompK*35, *ompK*36, GD or TD insertions, or combinations thereof, highlighting porin remodelling as a conserved complementary resistance mechanism. Lineage-stratified analysis revealed non-random carbapenemase distributions, with ST147 enriched for *bla*_NDM_ variants and ST231 predominantly associated with *bla*_OXA-232_, while *bla*_CTX-M-15_ showed no lineage restriction.

### 3.4 Genomic characterization of virulence genes

Virulence profiling revealed marked lineage-specific variation in aerobactin carriage (*iucABCD–iutA*) among the dominant carbapenem-resistant *K. pneumoniae* clones. Aerobactin was highly prevalent in ST2096 (84.68%, n=94/111) and ST231 (72.22%, n=78/108), whereas substantially lower carriage was observed in ST147 (40.7%, n=64/157). Overall, 62.7% (n=236/376) of isolates carried the aerobactin operon. The salmochelin operon (*iro*) and genotoxin-producing gene (colibactin, *clb*) were absent in all, while the yersiniabactin locus (ybt) was identified in 85.6% (n=322/376), often associated with ICEKp. Hypermucoviscosity-associated loci were variably distributed, with *rmpADC* detected in 43/376 isolates, *rmpA2* in 157/376, and *peg-344* in 158/376. Notably, phenotypic hypermucoviscosity did not correlate directly with the presence of *rmpA/rmpA2* Lineage-level analysis showed strong association of *iuc*5 with ST231 in the study isolates, while ST147 and ST2096 were associated with *iuc*1. The co-occurrence of three virulence markers, *rmpA/rmpA2,* with aerobactin was most evident in ST147 and ST2096, whereas ST231 typically carried aerobactin in the absence of these regulators.

### 3.5 Phylogenetic Analysis

Phylogenetic analysis based on the pan genome–derived core genome alignment revealed a strongly structured population composed of three deeply divergent clades corresponding to ST147, ST231 and ST2096 (**Figure S1**). These clades were mutually exclusive, with no evidence of phylogenetic intermixing between sequence types, consistent with long standing, lineage restricted evolutionary trajectories rather than recent cross lineage diversification. Study isolates were placed within their respective ST[specific clades with short terminal branches, indicating recent clonal expansion within each lineage.

Within each sequence type, the presence of multiple well-defined subclades (BAPS clusters) indicated ongoing diversification and clonal expansion. The presence of virulence-associated loci within discrete subclades is consistent with independent acquisition events followed by clonal expansion within each lineage. Notably, these virulence loci were recurrently associated with conserved capsule types and plasmid backgrounds, reinforcing the role of stable lineage-specific genetic contexts in shaping antimicrobial resistance and virulence convergence.

#### 3.5.1. Phylogenetic structure and virulence acquisition in ST147

Core-genome phylogenetic analysis of 444 *K. pneumoniae* ST147 genomes, including 157 study isolates, revealed notable intra-clonal diversity within this extensively disseminated high-risk clone (**Figure 1**). A maximum-likelihood tree, inferred from 629 core genes, bifurcated the phylogeny into two major clades, designated Clade 1 and Clade 2. Clade 1 exhibited notable genetic heterogeneity and was further subdivided into six BAPS clusters (1.1–1.6). In contrast, Clade 2 showed comparatively limited diversity and comprised two BAPS clusters (2.1 and 2.2), indicating more constrained evolutionary diversification. Study isolates displayed a highly non-random phylogenetic distribution, with approximately 90% clustering within three dominant sub-lineages (BAPS clusters 1.2, 1.6, and 2.1). The presence of two major phylogenetic branches, along with monophyletic clustering of study isolates within globally distributed genomes, supports a structured and non-random population organization within the ST147 lineage. Capsular locus analysis revealed marked heterogeneity across the ST147 phylogeny, with approximately 15 distinct K-loci identified, representing the highest capsule diversity observed among the analyzed lineages (**Figure 1**). Although KL64, KL51, and KL10 predominated, several additional capsule types were present at lower numbers and showed strong correspondence with phylogenetic clustering, indicating lineage-associated capsule diversification. Notably, virulence locus–positive ST147 isolates were predominantly associated with the KL64 capsular background, whereas other K-loci were more frequently observed among virulence-negative clusters.

**Figure 1.**
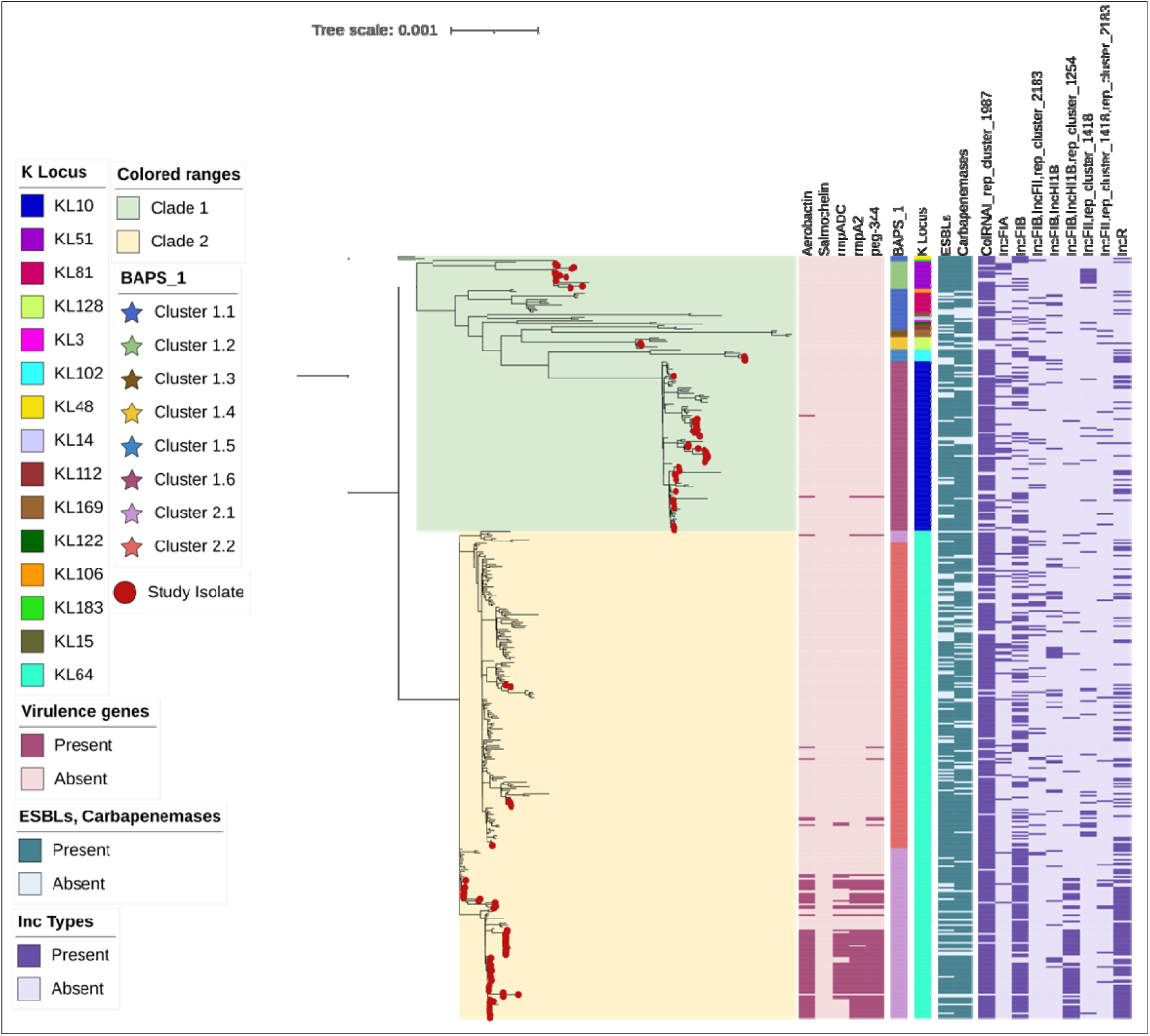
Midpoint-rooted maximum-likelihood core-genome phylogenetic tree of [n=444] *K. pneumoniae* ST147 isolates from India (n=157, marked with red circular symbols at terminal branches) and global sources, constructed from recombination-free core SNPs using IQ-TREE 2 (http://www.iqtree.org/) with 10,000 ultrafast bootstrap replicates. Two major sub-clades and eight clusters inferred using Bayesian Analysis of Population Structure (clusters); study isolates cluster prominently and are highlighted with distinct colours (clusters 1.6 and 2.1; Tapestry and Light Wisteria). Annotated metadata tracks from inner to outer illustrate (i) virulence genes heatmap; (ii) BAPS clusters; (iii) K-locus types, with KL64 and KL10 predominating among study isolates alongside 13 additional less frequent capsule loci (e.g., KL107, KL112); (iv) presence/absence of extended-spectrum β-lactamase (ESBL) genes (e.g., *bla*_CTX-M-15_ widespread); (v) plasmid replicon types heatmap; and (vi) carbapenemase genes (e.g., *bla*_NDM-5_). The figure was generated and annotated using iTOL (https://itol.embl.de/). Branch lengths represent nucleotide substitutions per site, with the scale bar (0.001 substitutions/site) indicating evolutionary divergence.

Virulence acquisition within ST147 was strongly cluster-restricted. Hybrid resistance–virulence plasmids were confined almost exclusively to cluster 2.1 and were absent from basal nodes across the phylogeny. In contrast, carbapenem resistance determinants were widely distributed across the ST147 phylogeny, with multiple *bla*_NDM_ variants (NDM-1 and NDM-5), *bla*_OXA-48-like_ variants (OXA-181/OXA-232) and ESBLs, predominantly *bla*_CTX-M-15,_ detected across both major clades and multiple BAPS clusters. Within cluster 2.1, isolates harbored IncFIB/IncHI1B/Rep hybrid plasmids carrying multiple antimicrobial resistance determinants (including *sul1/sul2*, *dfrA*, *mphE*, and *mrsE*) alongside key virulence loci (*iuc, rmpADC, rmpA2*, and *peg-344*); the salmochelin biosynthesis gene *iroB*, which is typically associated with virulence plasmids, was uniformly absent. Variation in virulence gene content and plasmid architecture among closely related isolates within this cluster suggests multiple independent acquisition events and/or ongoing plasmid remodelling rather than a single ancestral acquisition. The confinement of virulence loci to derived terminal subclades, together with their absence in basal backgrounds and in other clusters, indicates recent plasmid-mediated acquisition followed by clonal expansion. Although resistance–virulence hybrids were rare among global ST147 genomes, their high frequency among Indian study isolates suggests that local hospital-associated selective pressures favor plasmid acquisition and retention. Collectively, the increasing complexity of plasmid replicon content in virulent terminal branches highlights the central role of mobile genetic elements in driving rapid adaptive evolution within ST147 in clinical settings.

#### 3.5.2. Phylogenetic structure and virulence in ST2096

In contrast to the dynamic, cluster-restricted virulence acquisition observed in ST147, core-genome phylogenetic analysis of 259 *K. pneumoniae* ST2096 genomes revealed a highly conserved lineage with limited phylogenetic diversification (Figure 2). Study isolates clustered almost exclusively within BAPS clusters 2, 3, and 4, with no representatives in cluster 1, indicating a narrow population structure consistent with recent clonal expansion rather than deep lineage divergence. All genomes uniformly carried the KL64 capsular locus, further underscoring the genetic homogeneity of this lineage.

**Figure 2.**
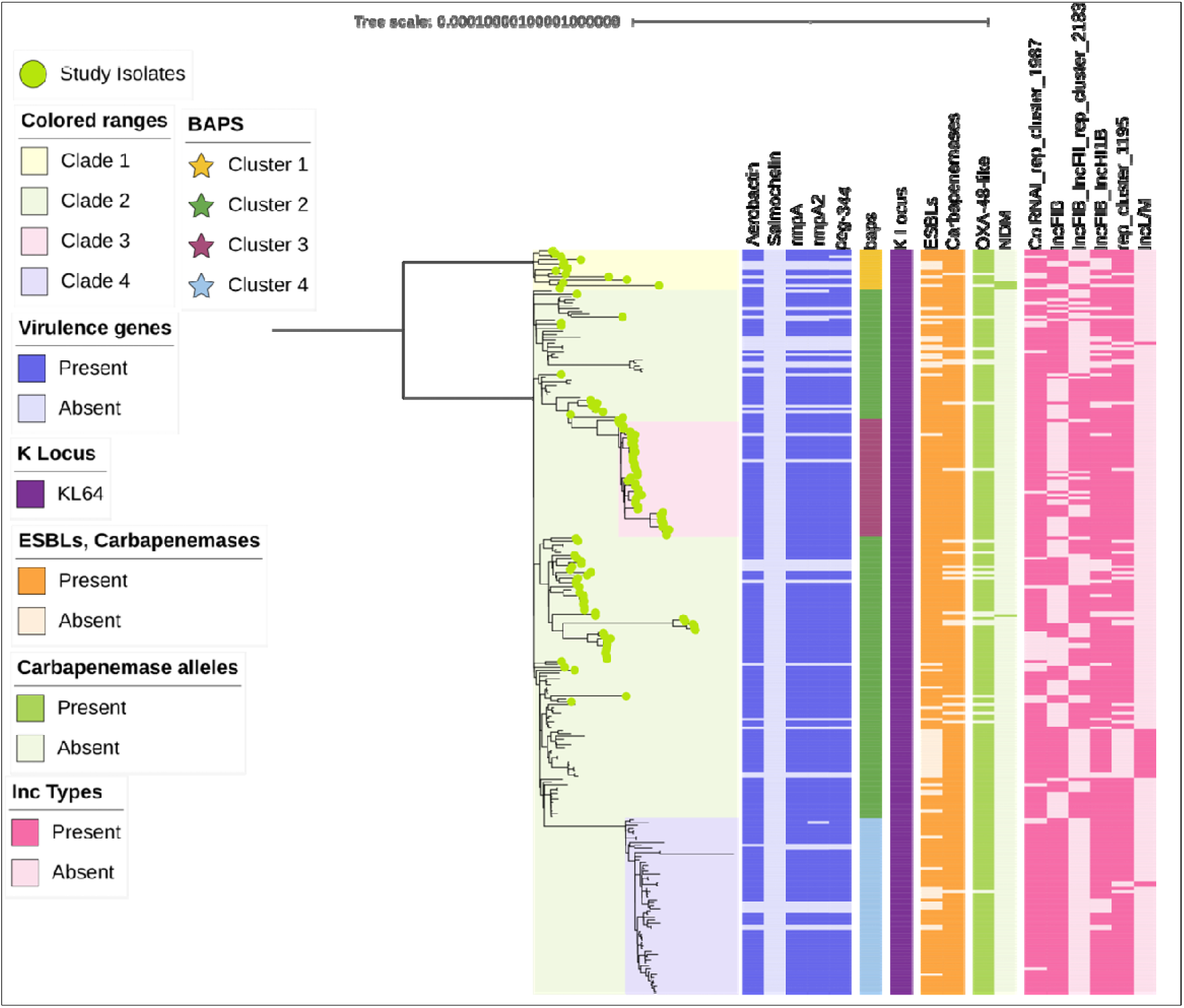
Midpoint-rooted maximum-likelihood core-genome phylogenetic tree of [n=259] *K pneumoniae* ST2096 isolates from India (n=111, marked with light green circular symbols at terminal branches) and global sources, constructed from recombination-free core SNPs using IQ-TREE 2 (http://www.iqtree.org/) with 10,000 ultrafast bootstrap replicates. Major sub-lineages inferred using BAPS are indicated as distinct clusters (Clusters 1–4: colours-Saffron, Asparagus, Cadillac and Cornflower), highlighted by colored star symbols; study isolates enrich Cluster 1 (dark green). Annotated metadata tracks from inner to outer depict (i) key virulence determinants including *peg-344* (prevalent in select clusters), visualized as heatmap; (ii) BAPS clusters; (iii) capsule locus types, with KL64 predominating near-universally; (iv) presence/absence of ESBL genes; (v) carbapenemase genes (e.g., *bla*_OXA-181_); and (vi) plasmid replicon content (e.g., IncF, ColRNAI variable). The figure was generated and annotated using iTOL (https://itol.embl.de/). Branch lengths represent nucleotide substitutions per site, with the scale bar providing a reference for evolutionary divergence.

**Figure 3:**
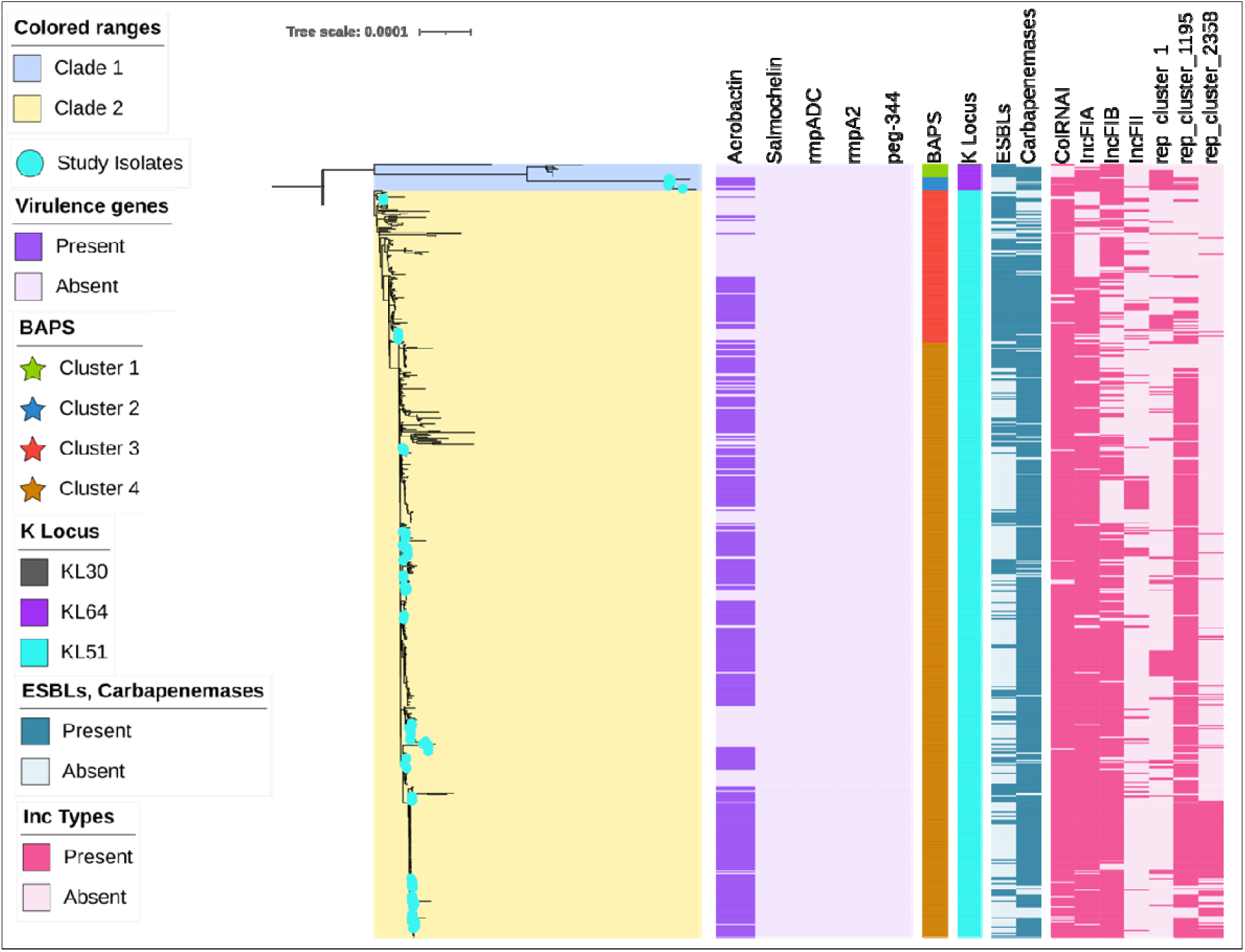
Phylogenetic relationships of ST231 Kp from India and global isolates Midpoint-rooted maximum-likelihood core-genome phylogenetic tree of [n=616] Kp ST231 isolates from India (n= 108, marked with teal circular symbols at terminal branches) and global sources, constructed from recombination-free core SNPs using IQ-TREE 2 (http://www.iqtree.org/) with 10,000 ultrafast bootstrap replicates. Major clusters inferred using BAPS are indicated as (Clusters 1–4; colours: Pistachio, Curious Blue, Red Orange and Meteor), highlighted by coloured star symbols; study isolates spread overall [teal]. Annotated metadata tracks from inner to outer depict (i) key virulence determinants including aerobactin (iuc operon), visualized as heatmaps; (ii) BAPS clusters; (iii) capsule locus types, with KL51 predominating; (iv) plasmid replicon content (e.g., IncFIA, IncFIB); (v) presence/absence of ESBL genes (e.g., *bla*_CTX-M-15_); (vi) carbapenemase genes (e.g., *bla*_OXA-232_). The figure was generated and annotated using iTOL (https://itol.embl.de/). Branch lengths represent nucleotide substitutions per site, with the scale bar (0.001 substitutions/site) providing a reference for evolutionary divergence.

Unlike ST147, virulence determinants including the aerobactin locus (*iuc*), *peg-344*, and *rmpA2*, were distributed broadly across the phylogeny, including basal nodes, suggesting early acquisition followed by long-term stable maintenance. Although *rmpA* was detected sporadically, the remaining *rmpDC* genes were consistently absent, and similar to ST147 salmochelin biosynthesis locus (*iro*) was uniformly lacking across the lineage. Aerobactin carriage showed a strong and conserved association with integrated IncFIB and IncHI1B replicons across all BAPS clusters, indicating that plasmid-mediated virulence acquisition has subsequently stabilized rather than undergone ongoing gain or loss.

In addition to virulence loci, this IncFIB–IncHI1B-associated plasmid frequently co-harbored accessory resistance genes, including tet, *bla*_TEM_, *bla*_CTX-M-15_, *aadA*, *dfrA*, and *sul*, reflecting a stable linkage between aerobactin-associated virulence and non-carbapenem resistance traits. Resistance profiles further mirrored this phylogenetic stability. ST2096 was strongly associated with *bla*_OXA-48-like_ carbapenemases, predominantly *bla*_OXA-232_, typically carried on small Col-type plasmids that co-existed with the integrated virulence plasmid and were consistently maintained across the lineage. Collectively, the conserved maintenance of capsular type, virulence loci, resistance determinants, and plasmid replicon composition indicates that ST2096 represents a stabilized resistance–virulence background, in stark contrast to the ongoing plasmid-driven diversification observed in ST147.

#### 3.5.3. Phylogenetic structure and virulence in ST231

Core-genome phylogenetic analysis of *K. pneumoniae* ST231 revealed clear phylogenetic diversity, with a small number of divergent isolates forming a basal phylogenetic cluster and the majority of genomes comprising a closely related backbone cluster with short internal branches, consistent with recent clonal expansion. Study isolates were distributed across both the divergent basal cluster and the dominant backbone cluster, indicating that the dataset captures both ancestral diversity and contemporary circulating lineages. Capsular locus variation mapped closely to phylogenetic position. The divergent basal isolates were associated with alternative K-loci, including KL64 and KL30, whereas the expanding backbone was uniformly characterized by the KL51 capsular locus. Although BAPS analysis subdivided this backbone into multiple clusters, these subclusters within backbone cluster showed minimal phylogenetic separation and shared the same capsular background, suggesting fine-scale population structure within a single expanding KL51 lineage rather than biologically distinct capsule-associated lineages. Antimicrobial resistance in ST231 was characterized by a high prevalence of ESBLs, predominantly *bla*_CTX-M-15_, alongside carbapenem resistance mediated mainly by *bla*_OXA-232_. In contrast to ST147 and ST2096, canonical hypervirulence-associated markers, including *peg-344*, *rmpA*, and *rmpA2*, were uniformly absent across all ST231 isolates. Virulence determinants detected in ST231 were limited primarily to the aerobactin biosynthesis operon (iuc), which was present across both basal and backbone lineages, consistent with early acquisition followed by stable maintenance.

Plasmid replicon analysis revealed a highly conserved plasmid architecture across ST231 phylogeny. The iuc operon was predominantly associated with IncFIA plasmids, while carbapenem resistance determinants were maintained on distinct plasmid replicons. In addition to iuc, IncFIA plasmids frequently co-harbored accessory antimicrobial resistance genes, including *erm*, *mphA*, *aadA2*, and *dfrA12*, along with the class 1 integron integrase gene intI1. This segregation of virulence and resistance elements was conserved across phylogenetically divergent and derived clusters and showed no correspondence with phylogenetic subdivision, indicating lineage-embedded and evolutionarily stable plasmid configurations rather than ongoing plasmid flux. Together, these findings position ST231 as an intermediate evolutionary state between the highly dynamic plasmid-driven diversification observed in ST147 and the stabilized resistance–virulence convergence characteristic of ST2096.

## 4. Replicon distribution and lineage-specific plasmids

Comparative plasmidome analysis revealed that carriage of multiple plasmid replicons was common across all three high-risk lineages, consistent with the plasmid-rich genomic background of *K. pneumoniae*. However, marked lineage-specific differences were observed in the organization and integration of virulence-associated plasmids. In ST147, replicons associated with antimicrobial resistance were highly diverse, with ColKP3 (62%), IncFII (47%), and IncR (41%) predominating, followed by IncFIB variants and IncHI1B(pNDM-MAR) (∼34%). Virulence-associated markers from phylogenetic analysis were predominantly linked to large co-integrated IncFIB–IncHI1B replicons (∼300 kb). In contrast, ST2096 exhibited a conserved plasmid replicon profile despite frequent multi-replicon carriage. Nearly all isolates harbored IncFIB(K)_1_Kpn3 (94%), IncFIB(pNDM-MAR) (86%), and IncHI1B(pNDM-MAR) (84%), with 82% carrying hybrid IncFIB-IncHI1B plasmids (∼250–350 kb) as the dominant hybrid inc types preserved across isolates. ST231 displayed a distinct plasmid profile. ColKP3 replicons (72%), typically carrying *bla*_OXA-232_, together with IncFIA (60%) and IncFII (52%), predominated. Notably, IncFIB–IncHI1B hybrid replicons were absent. Virulence-associated markers (only iuc here) were instead localized to smaller IncFIA plasmids (∼70 kb), suggesting an alternative mode of virulence carriage relative to ST147 and ST2096.

## 5. Lineage-Specific Resistance–Virulence Plasmid Architectures

To investigate the structural basis of resistance–virulence convergence, representative isolates from ST147 (n = 15), ST2096 (n = 15), and ST231 (n = 5) underwent long-read ONT sequencing and plasmid reconstruction. A smaller number of ST231 isolates was included because preliminary analysis showed highly conserved plasmid structures with limited variation across the lineage. Comparative plasmid analysis identified three distinct lineage-specific plasmid architectures associated with resistance–virulence convergence (Figures 4–6).

**Figure 4.**
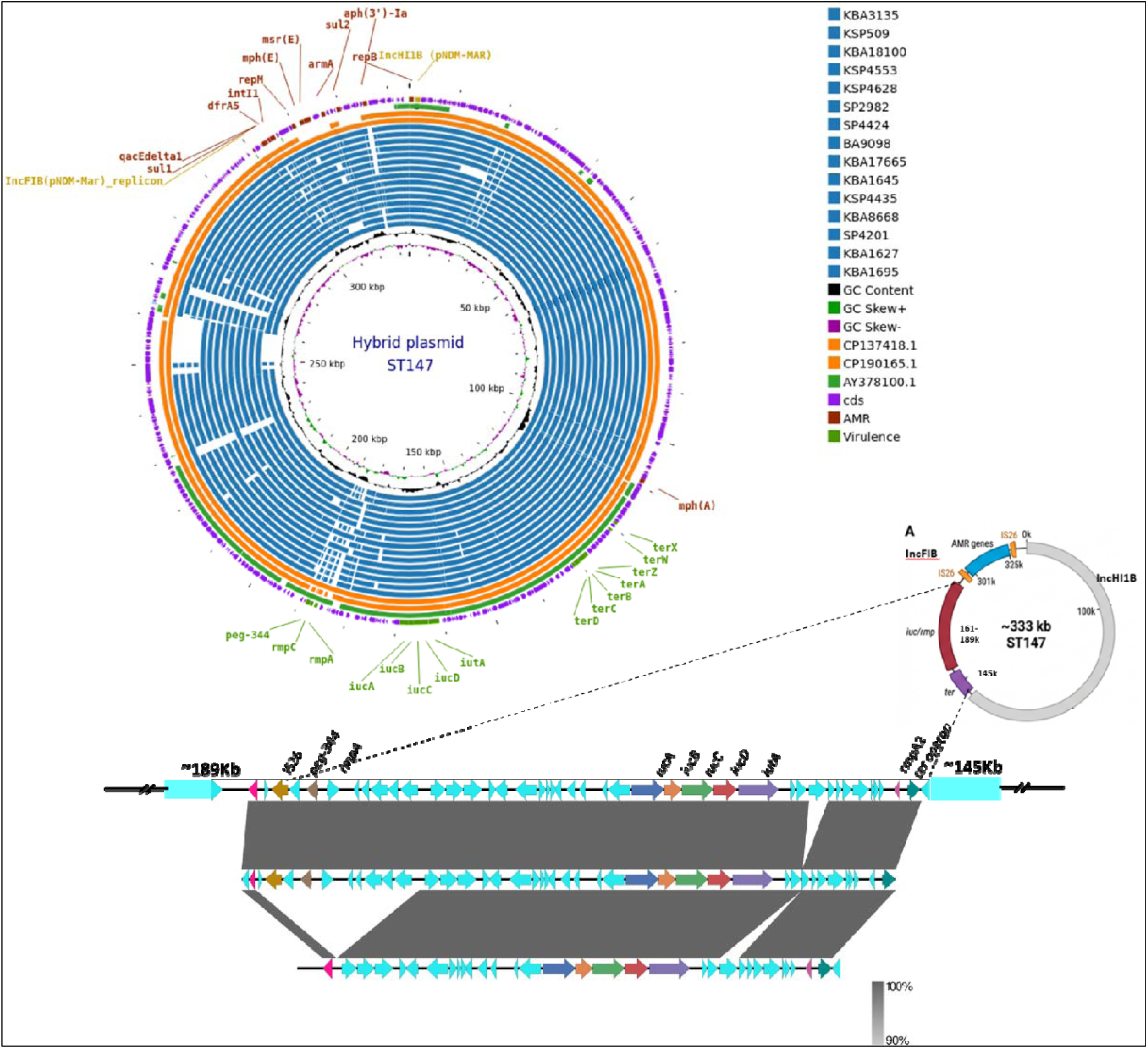
Comparative structural analysis of IncHI1B–IncFIB hybrid virulence–resistance plasmids identified in ST147 isolates. (A) Circular alignment of representative hybrid plasmids from ST147 isolates in this study (n = 15) compared with reference plasmids retrieved from GenBank (CP137418.1, CP190165.1, and AY378100.1). Rings from outer to inner circles represent plasmids from the study isolates followed by the reference plasmids. Conserved backbone regions, virulence-associated loci (*iucABCD-iutA, rmpA/rmpA2, peg-344*), antimicrobial resistance determinants, GC content (black), and GC skew (green and magenta) are shown. (B) Schematic representation of the representative ∼333 kb ST147 hybrid plasmid highlighting the IncFIB and IncHI1B replicons, virulence region (*iuc/rmp* locus), antimicrobial resistance module, and tellurite resistance (*ter*) operon. (C) Linear synteny comparison of representative hybrid plasmids generated using EasyFig, demonstrating conserved backbone architecture with variable insertion and rearrangement regions associated with virulence and antimicrobial resistance modules.

**Figure 5:**
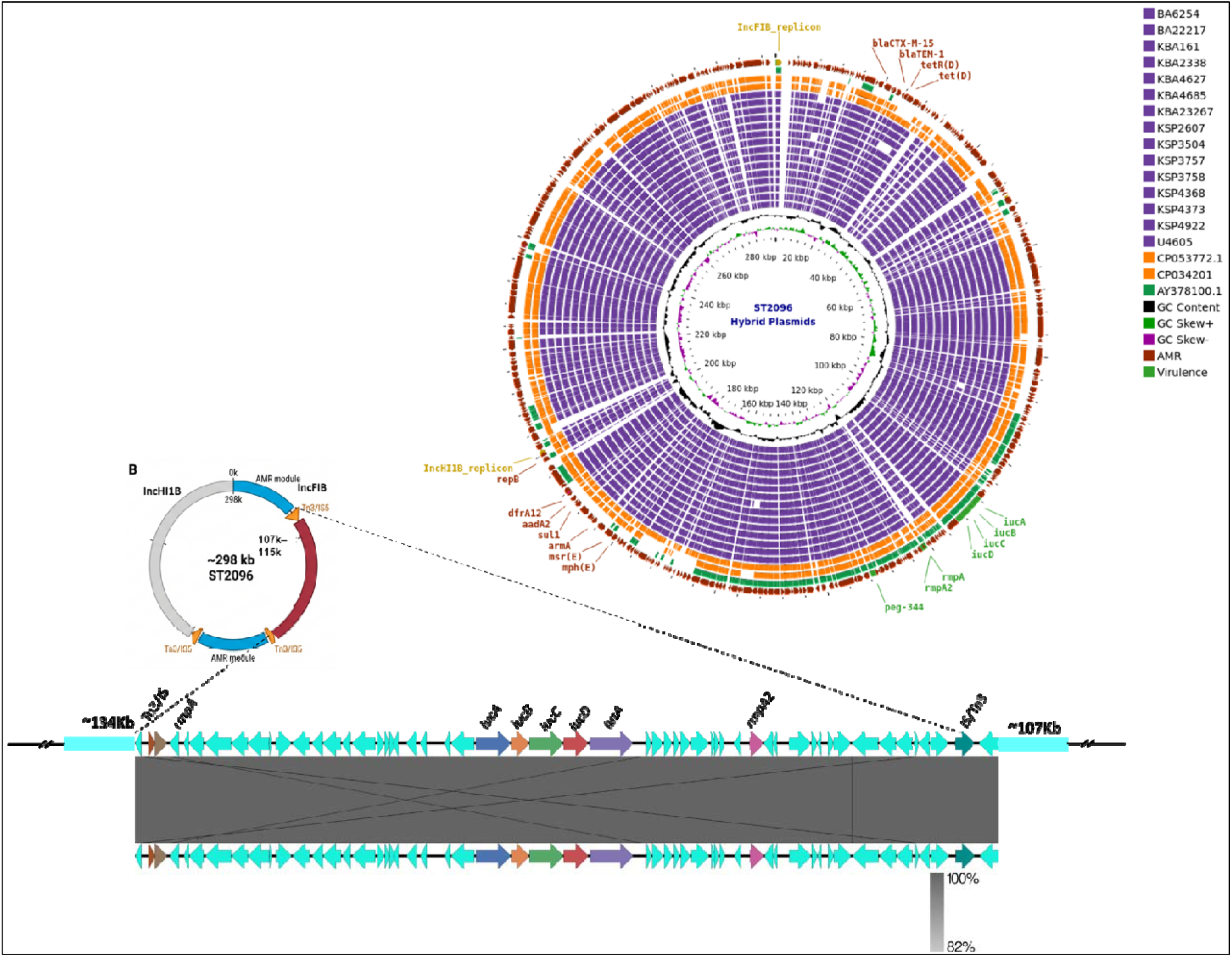
Comparative structural analysis of IncHI1B–IncFIB hybrid virulence–resistance plasmids identified in ST2096 isolates. (A) Circular alignment of representative hybrid plasmids from ST2096 isolates in this study (n = 15) compared with previously reported reference plasmids retrieved from GenBank (CP053772.1, CP034201.1, and AY378100.1). Rings from outer to inner circles represent plasmids from the study isolates followed by the reference plasmids. Conserved backbone regions, virulence-associated loci (*iucABCD-iutA, rmpA/rmpA2, peg-344*), antimicrobial resistance determinants, GC content (black), and GC skew (green and magenta) are shown. (B) Linear synteny comparison of representative ST2096 hybrid plasmids generated using EasyFig, illustrating the conserved plasmid backbone with variable insertion and rearrangement regions associated with virulence and antimicrobial resistance modules.

**Figure 6:**
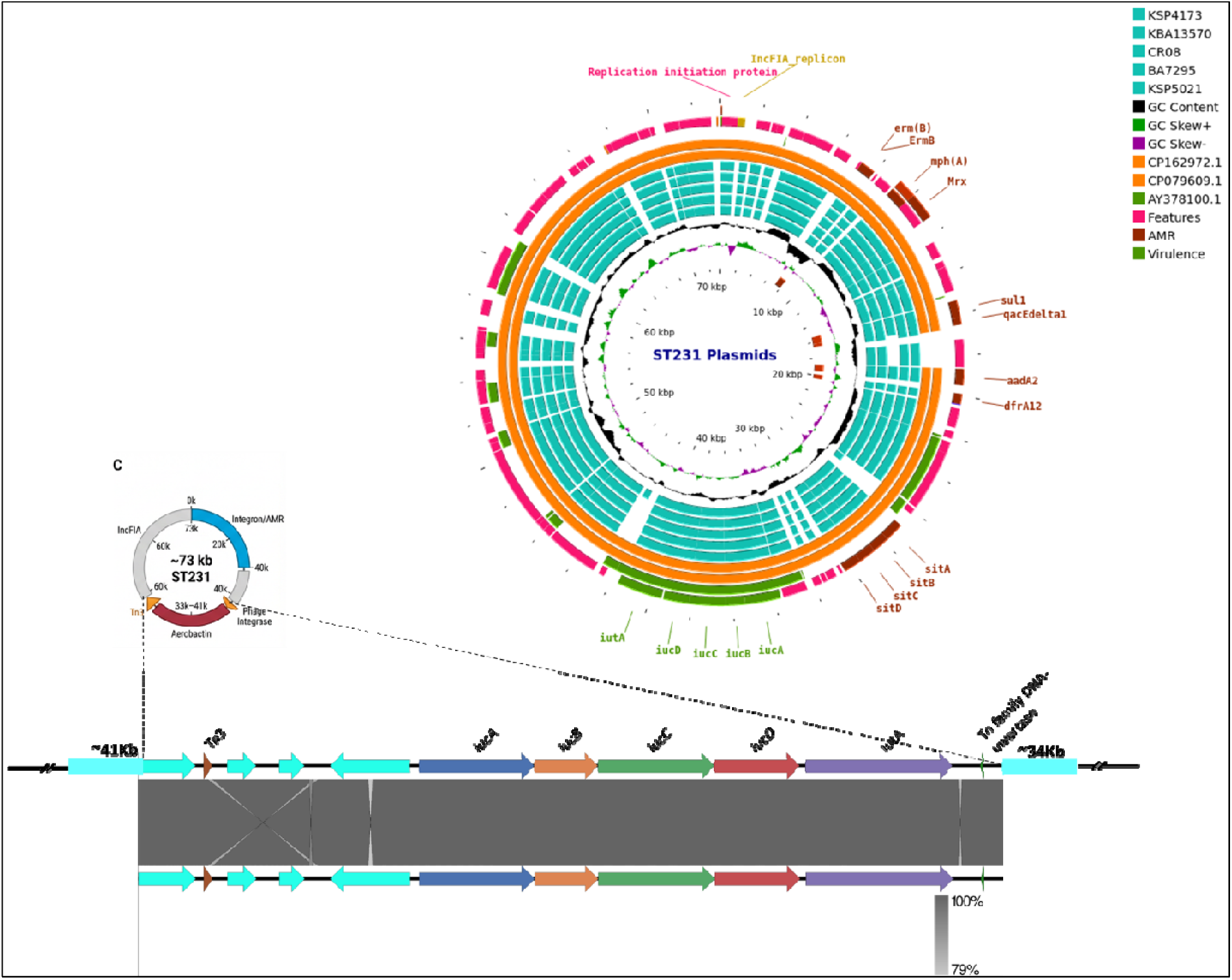
Comparative structural analysis of virulence-associated plasmids identified in ST231 isolates. (A) Circular maps of representative plasmids from ST231 isolates in this study (n = 5) compared with reference plasmids retrieved from GenBank (CP162972.1, CP079609.1, and AY378100.1). The analysis demonstrates the presence of IncFIA-associated virulence plasmids carrying the aerobactin locus (*iucABCD-iutA*) and distinct ColKP3 plasmids harbouring *bla*_OXA-232_, shown separately. Annotated features include plasmid replication regions, virulence-associated loci, antimicrobial resistance determinants, GC content (black), and GC skew (green and magenta). (B) Linear synteny comparison of representative ST231 plasmids generated using EasyFig, illustrating conserved plasmid architecture and homologous regions across virulence-associated and resistance plasmids.

None of the major lineages carried complete classical hypervirulence plasmids typically associated with community-acquired hvKp. Instead, all lineages showed partial acquisition of virulence-associated loci within multidrug-resistant hospital-adapted backgrounds, consistent with an intermediate convergence state between classical MDR-Kp and fully hypervirulent CR-hvKp.

In ST147, virulence determinants including the aerobactin locus (*iucABCD–iutA*) together with variable *rmpA/rmpC* and *peg-344* were integrated within large IncHI1B–IncFIB hybrid plasmids carrying multiple antimicrobial resistance genes. These plasmids also contained several mobile genetic elements, suggesting ongoing plasmid remodeling and structural diversity. Extensive backbone regions were additionally observed in full-length plasmid comparisons, supporting active plasmid variations within the ST147 lineage (Figure S2).

In contrast, ST2096 displayed highly conserved IncFIB–IncHI1B hybrid plasmids with stable organization of both virulence and resistance regions across isolates. Full-length plasmid comparisons against the reference showed structural conservation across ST2096 plasmids, with only limited insertion-associated no variation observed between isolates (Figure S3). These plasmids consistently carried *iucABCD–iutA, peg-344,* and *rmpA2,* while carbapenemase genes (*bla*_NDM-5_/*bla*_OXA-232_) were usually located on separate plasmids.

ST231 showed a different plasmid organization in which virulence and resistance determinants remained largely segregated. The aerobactin locus was mainly carried on IncFIA virulence plasmids lacking classical hypervirulence markers such as *rmpA2* and *peg-344*, whereas carbapenem resistance was mediated separately through ColKP3 plasmids carrying *bla*_OXA-232_. Unlike ST147 and ST2096, IncFIB–IncHI1B hybrid plasmids were not identified in ST231. Comparative full-plasmid representation further confirmed the presence of the virulence-associated region on distinct IncFIA-associated plasmids in ST231 isolates (Figure S4). Together, these findings demonstrate that resistance–virulence convergence among dominant Indian hospital-associated *K. pneumoniae* lineages occurs through distinct lineage-specific plasmid configurations rather than dissemination of a single convergent plasmid type.

## 4. Discussion

The coexistence of both multidrug resistance genes and virulence determinants within the same bacterial background represents a major evolutionary shift in hospital-associated *K. pneumoniae*. Globally, the emergence of these resistance–virulence convergent high-risk clones has largely been explained through two classical routes (36). The first is acquisition of MDR plasmids by highly virulent ST23-like lineages. The second is transfer of pLVPK-like virulence plasmids into established multidrug-resistant clones, particularly ST11 (37, 38). The latter mechanism has been reported most frequently in South and East Asia where it has driven several outbreaks of carbapenem-resistant hypervirulent *K. pneumoniae* (27, 41). However, resistance–virulence convergence is an ongoing and complex evolutionary process, and different lineages may follow distinct pathways to reach similar convergent states. Our findings support the view that acquisition and long-term maintenance of complete virulence plasmids by multidrug-resistant clones may be relatively uncommon. Classical pLVPK-like virulence plasmids are largely restricted to specific hypervirulent genetic backgrounds and appear difficult to disseminate widely among multidrug-resistant lineages because they are typically non-self-transmissible and may impose substantial fitness costs in unrelated hosts (39). Consequently, successful convergence often depends on additional evolutionary mechanisms that facilitate the transfer, maintenance, and stabilization of virulence determinants within resistant lineages (40). In this regard, recent studies have highlighted the importance of homologous recombination, helper plasmids, and formation of resistance–virulence hybrid plasmids in driving convergence. These observations emphasize that the evolutionary routes leading to MDR-hvKp are considerably more diverse than initially predicted (33).

Against this background, our findings suggest that dominant Indian hospital-associated clones have adopted a distinct convergence strategy. Rather than acquiring complete virulence plasmids, these lineages selectively retain specific virulence determinants through diverse resistance–virulence plasmid architectures. Across ST147, ST2096, and ST231, the aerobactin locus was consistently present, whereas other canonical hypervirulence-associated determinants including *iro, rmpA,* and *rmpA2* showed variable distribution or were absent altogether, defining a shared intermediate convergence state across all three lineages (30, 51). The consistent retention of aerobactin across three independent high-risk lineages suggests that it provides a selective advantage. As a major virulence determinant, aerobactin plays a key role in iron acquisition, extraintestinal dissemination, and host adaptation (35, 49, 51). In contrast, maintenance of complete virulence plasmids and highly mucoviscous phenotypes, may impose considerable metabolic costs particularly when combined with large multidrug-resistance plasmids (39). Consistent with this hypothesis, convergent CR-hvKP lineages frequently show attenuation of hypermucoviscosity and reduced expression of *rmpA/rmpA2*-associated virulence pathways (48,50). Consequently, convergent multidrug-resistant isolates carrying canonical virulence loci often exhibit lower-than-expected virulence phenotypes (1). Together, these observations support the view that the intermediate convergence state observed among dominant Indian hospital clones is consistent with a hospital-adapted evolutionary optimum, in which selective retention of high-value determinants such as aerobactin balances invasive potential with long-term persistence under sustained antimicrobial selection.

Within this framework, ST147 appears to represent the most dynamic route towards the shared intermediate convergence state. As one of the most globally disseminated high-risk MDR clones, ST147 has repeatedly demonstrated remarkable genomic plasticity, with a well-recognized capacity to acquire carbapenemases, ESBL determinants, and diverse mobile genetic elements in hospital settings (8, 19). Increasing evidence suggests that this success is closely linked to the ability of ST147 to repeatedly recruit and remodel accessory genetic elements, particularly IncFIB - IncHI1B hybrid plasmids carrying combinations of resistance and virulence determinants (5, 6, 21). Notably, these hybrid plasmids do not represent a single conserved virulence–resistance platform. Instead, studies from Europe and East Asia have revealed multiple related but structurally distinct IncHI1B–IncFIB backbones carrying different assortments of virulence loci, resistance genes, and mobile genetic elements (6, 52). The extensive rearrangement of insertion sequences, transposons, and resistance islands reported within ST147 populations further suggests that these plasmids are subject to continual recombination and remodeling rather than stable vertical inheritance (14). Recent evidence of ongoing in-host evolution and convergence within ST147 further reinforces the notion that this lineage remains evolutionarily active and highly adaptable (17). Consistent with this model, virulence determinants in our collection were confined to specific phylogenetic subclusters and associated with diverse IncHI1B–IncFIB hybrid plasmid architectures. Together, these observations support the view that ST147 functions as an opportunistic assembler of resistance and virulence modules, in which continual plasmid restructuring serves as the primary mechanism driving resistance–virulence convergence.

By contrast, ST2096 represents a fundamentally different convergence trajectory characterized by stabilization rather than continual remodeling. First documented in India as a multidrug-resistant hypervirulent lineage, ST2096 is consistently associated with highly conserved IncFIB–IncHI1B hybrid plasmids carrying aerobactin and *rmpA2* (27). Unlike the mosaic, repeatedly remodelled plasmid configurations observed in ST147, the ST2096 hybrid appears to function as a stable, integrated unit. The presence of virulence determinants, within basal phylogenetic nodes suggests an early convergence event followed by clonal expansion (27). Similar convergent ST2096 isolates have been reported from geographically distinct settings, supporting the notion that this resistance–virulence configuration is both stable and epidemiologically successful (15, 42, 43). This pattern aligns with emerging models showing that certain IncHI1B-based hybrid plasmids can achieve stable transmission within permissive genomic backgrounds (46, 47). Notably, carbapenemase genes in this cohort were carried on separate, smaller plasmids rather than integrated into the large hybrid structure, suggesting modular coexistence of both traits without full plasmid fusion (27). ST2096 therefore appears to represent a consolidator lineage, in which a favorable resistance–virulence configuration has been evolutionarily stabilized and subsequently amplified through clonal expansion.

A third convergence strategy is exemplified by ST231, where resistance and virulence traits coexist without the formation of a large hybrid resistance–virulence plasmid. ST231 is a major carbapenem-resistant lineage in India and is strongly associated with *bla*_OXA-48-like_, frequently carried on small ColKP3 plasmids (34, 44). Despite this conserved resistance background, virulence determinants in our collection were primarily associated with smaller IncFIA plasmids carrying the aerobactin locus. In contrast to ST147 and ST2096, large IncFIB–IncHI1B hybrid plasmids were absent, indicating that convergence can occur without physical integration of resistance and virulence determinants into a single plasmid backbone. The continued recovery of ST231 from diverse clinical and carriage settings further indicates that stable coexistence of compatible resistance and virulence plasmids can support long-term persistence and dissemination without extensive plasmid fusion (15, 45). ST231 therefore appears to represent a segregator strategy, in which resistance and virulence determinants remain physically separated while still achieving the same intermediate convergence state observed in ST147 and ST2096.

When considered together, these three lineages demonstrate that the intermediate convergence state in hospital-associated *K. pneumoniae* can arise through multiple evolutionary pathways. ST147 represents a highly dynamic plasmid-remodeling pathway, ST2096 a stabilized hybrid-plasmid pathway, and ST231 a segregated plasmid co-maintenance pathway. Despite these distinct trajectories, all three lineages converge on a similar biological endpoint, highlighting that clinically comparable convergent phenotypes can emerge through fundamentally different plasmid architectures. These findings also underscore the limitations of relying solely on hypermucoviscosity or single virulence markers such as aerobactin for surveillance, as isolates carrying diverse combinations of resistance and virulence determinants may lack classical hypervirulent phenotypes while still representing biologically significant intermediate forms within high-risk multidrug-resistant backgrounds. Although genomic surveillance identifying co-occurring resistance and virulence loci has substantially improved detection of convergent strains, plasmid-level resolution remains essential for understanding the acquisition, maintenance, and dissemination of these traits (31, 32). Multiplex PCR assays targeting combinations of carbapenemase genes and major virulence markers may provide an efficient first-line screening approach (4); however, lineage-specific hospital based genomic surveillance and plasmid reconstruction remain critical for tracking both clonal spread and ongoing plasmid evolution within high-risk *K. pneumoniae* lineages (53).

Nevertheless, several constraints in our study design must be noted. First, long-read sequencing and subsequent plasmid reconstruction were limited to representative isolates from each lineage; while this approach facilitated detailed structural comparisons, it may not capture the full spectrum of plasmid diversity within these clonal populations. Second, the study was based on a carefully selected subset of Indian clinical isolates from a single hospital rather than comprehensive sequencing of all circulating ST147, ST2096, and ST231 strains, which may limit resolution of lineage-wide evolutionary patterns. Third, functional assays including plasmid transfer kinetics, stability testing, and *in vivo* virulence models were not performed. Future studies integrating broader genomic surveillance with longitudinal sampling and experimental validation will be important to further define the evolutionary dynamics of resistance–virulence convergence in high-risk *K. pneumoniae* lineages.

## 5. Conclusion

This study concludes that genomic and plasmid analysis showed that hospital-adapted *K. pneumoniae* lineages acquire and maintain virulence through distinct, lineage-specific plasmid configurations. Divergent patterns of convergence were seen in ST147, ST2096, and ST231, which were distinguished by segregated, stable or integrated plasmids. Convergence is driven by multiple evolutionary pathways shaped by clone-specific genetic backgrounds. Moreover, the lack of complete virulence plasmids, the constant presence of aerobactin, and the evident discordance between virulence genotype and hypermucoviscosity phenotype affirm that these clones exist in an intermediate convergent state. This state indicates a continuous evolution under antimicrobial selection. Thus, epidemiological surveillance in healthcare environments should extend beyond resistance-focused genotyping to include essential virulence markers in order to effectively monitor and reduce the dissemination of these predominant high-risk lineages.

## Acknowledgements

The authors thank the Department of Clinical Microbiology, Christian Medical College and Hospital, Vellore, for providing us with all the necessary facilities to conduct our study. This work forms part of the Ph.D. thesis of S.M.K., submitted to The Tamil Nadu Dr. M.G.R. Medical University.

## Author Contributions

S.M.K.: conceptualization, data curation, methodology, investigation, formal analysis, visualization, writing – original draft, and writing – review and editing. J.J.J.: data curation, validation, and writing – review and editing. S.R.: methodology, software, and formal analysis. M.P.: formal analysis. K.G.: investigation and formal analysis. A.V.: formal analysis. A.N. and R.N.: resources and validation. A.M.: supervision and clinical oversight. R.E.: writing – review and editing. K.W.: project administration, funding acquisition, supervision, and writing – review and editing. B.V.: conceptualization, supervision, project administration, funding acquisition, and writing – review and editing.

## Ethical statement

The study was approved by the Institutional Review Board (IRB), Christian Medical College, Vellore, min. no. 15248, dated 22 March 2023.

