## Supplementary Figures 1-4 for "Three Plasmid Strategies, One Intermediate Convergence State: Lineage-Specific Resistance–Virulence Architecture in Dominant Indian Carbapenem-Resistant *Klebsiella pneumoniae* Clones": Supplementary Figures.docx

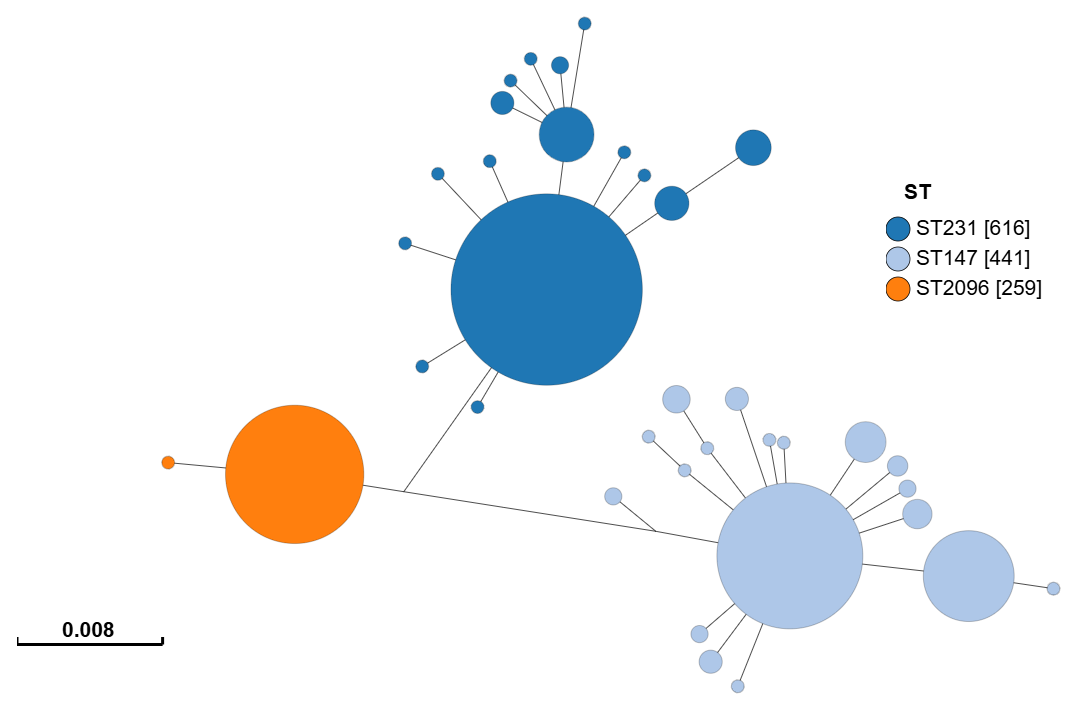
**Figure S1.** Minimum spanning tree based on cgMLST showing the global population structure of 1,400 *K pneumoniae* isolates belonging to high-risk sequence types ST231, ST147, and ST2096.

The tree was constructed using allelic profiles derived from the cgMLST scheme, and visualized using GrapeTree. Each node represents a unique cgMLST allelic profile, and node size is proportional to the number of isolates sharing that profile. Nodes are coloured according to sequence type (ST231, dark blue; ST147, light blue; ST2096, orange). Edges connecting nodes represent allelic differences between profiles, and edge lengths are proportional to the genetic distance, with the scale bar indicating the number of allelic substitutions per locus. Clustering patterns highlight the genetic relatedness, lineage structure, and diversification within and between major high-risk clones. The tree includes both study isolates and globally distributed reference genomes, providing phylogenetic context and illustrating lineage expansion and divergence. This analysis enables visualization of clonal relationships and evolutionary structure at the population level.

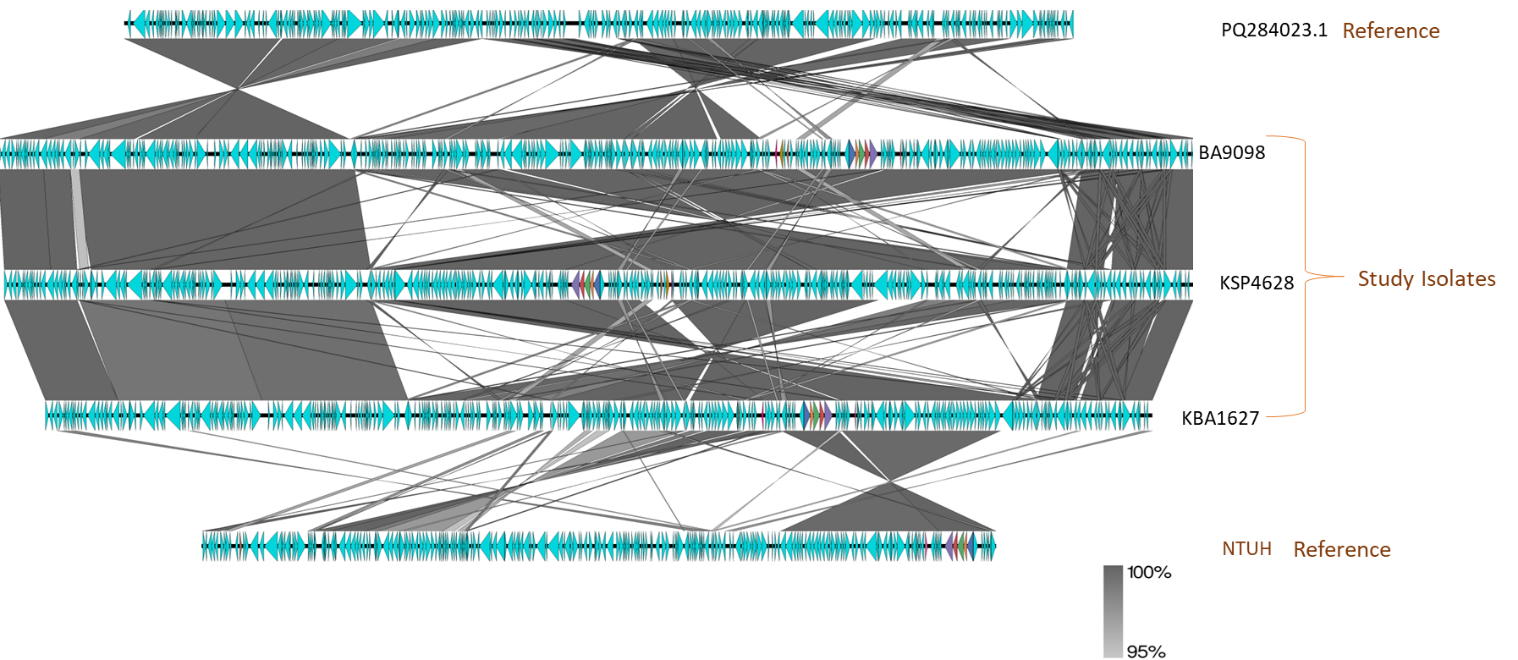

**Figure S2.** Comparative full-length plasmid alignment of representative ST147 hybrid plasmids generated using EasyFig. The uppermost plasmid (PQ284023.1) represents a reference IncHI1B–IncFIB backbone lacking virulence-associated loci and was used to trace structural conservation across plasmids. Representative plasmids from study isolates (BA9098, KSP4628, and KBA1627) demonstrate insertion of virulence-associated regions containing *iuc-, rmp-,* and *peg-344*-associated loci while maintaining overall backbone conservation. The bottom plasmid (NTUH) represents a reference virulence plasmid used for comparison of the acquired virulence region. Shaded regions indicate homologous sequences shared between plasmids, highlighting lineage-specific rearrangements and insertion-associated plasmid remodeling.

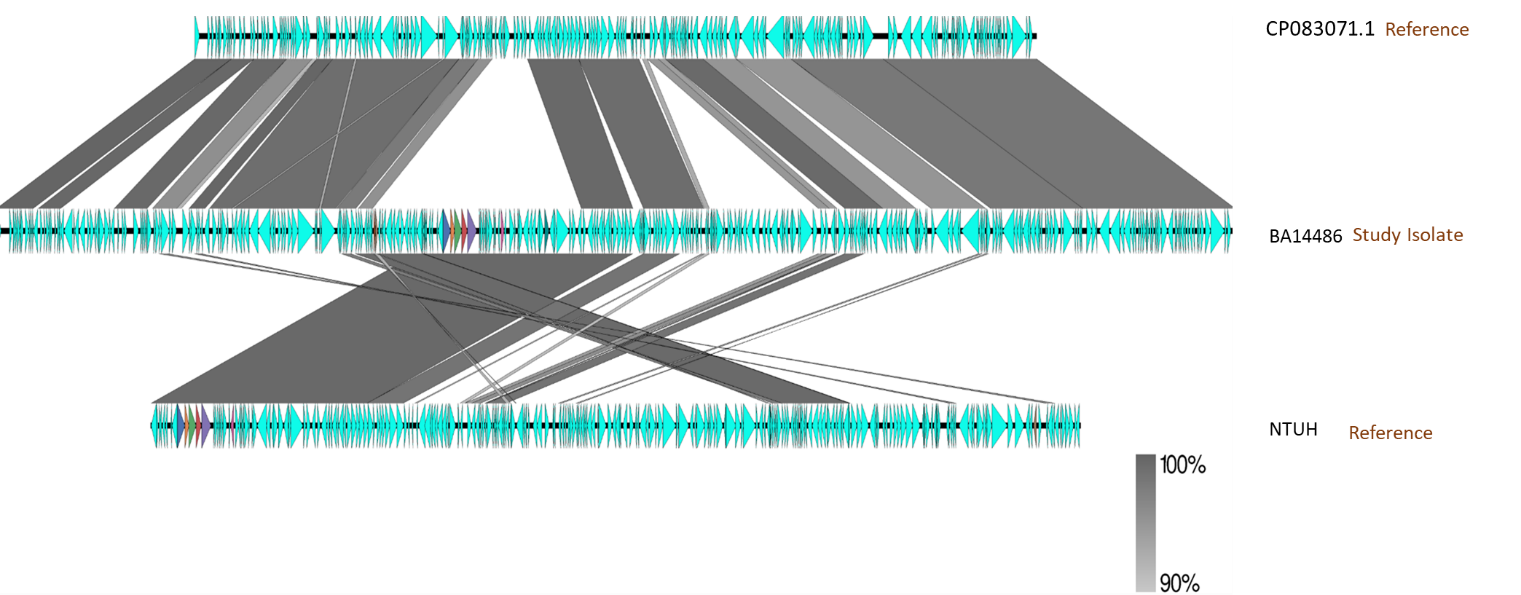

**Figure S3.** Comparative full-length plasmid alignment of representative ST2096 hybrid plasmids generated using EasyFig. The uppermost plasmid (CP083071.1) represents a reference IncHI1B–IncFIB backbone lacking virulence-associated loci and was used to assess structural conservation across plasmids. Representative plasmids from study isolates (BA14486) demonstrate acquisition of virulence-associated regions containing *iuc-, rmpA2-*, and *peg-344*-associated loci while preserving the conserved plasmid backbone. The bottom plasmid (NTUH) represents a reference virulence plasmid used for comparison of the acquired virulence region. Shaded regions indicate homologous sequences shared between plasmids, demonstrating high structural conservation within the ST2096 lineage.

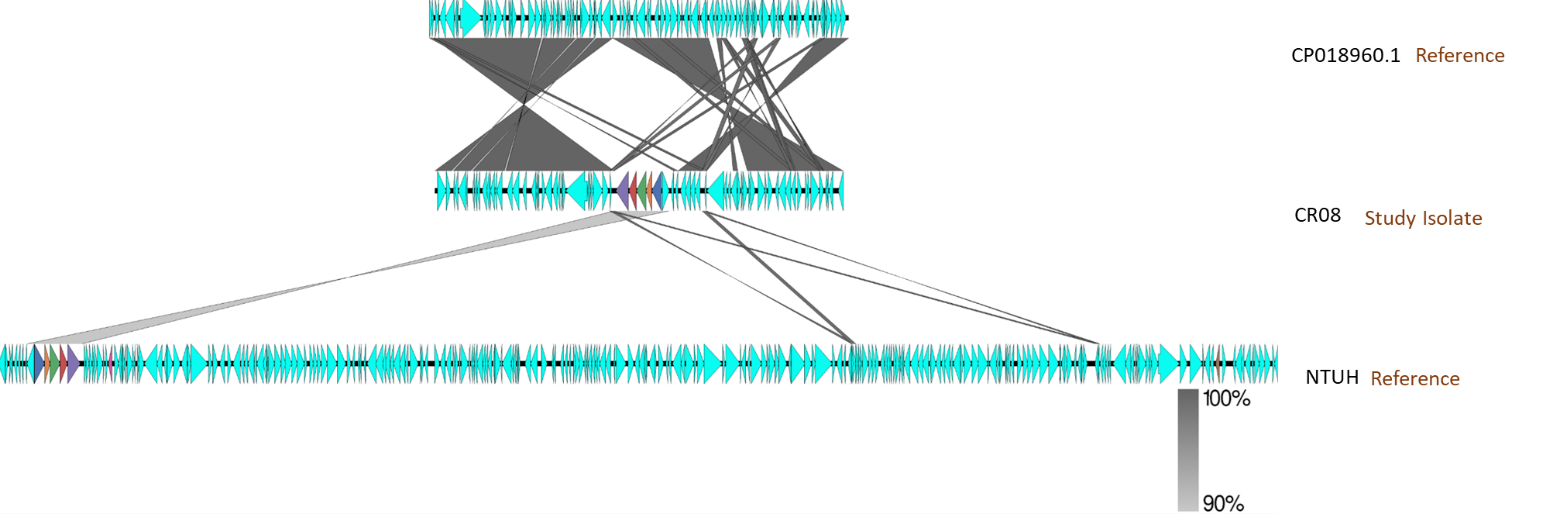

**Figure S3.** Comparative full-length plasmid alignment of representative ST231 virulence-associated plasmids generated using EasyFig. The uppermost plasmid (CP018960.1) represents a reference plasmid backbone lacking virulence-associated loci. Representative plasmids from study isolate CR08 demonstrate insertion of the aerobactin-associated virulence region while maintaining backbone conservation. The bottom plasmid (NTUH) represents a reference virulence plasmid used for comparison of the acquired virulence-associated region. Shaded regions indicate homologous sequences shared between plasmids, confirming localization of virulence-associated loci on distinct IncFIA-associated plasmid backbones in ST231 isolates.
